# Mesenchymal WNT2B is required for the development and function of the human intestine

**DOI:** 10.1101/2025.05.01.651727

**Authors:** Akaljot Singh, Holly M. Poling, Jennifer Foulke-Abel, Nambirajan Sundaram, Abid A. Reza, Sarah Joseph, Abrahim ElSeht, Kalpana Srivasta, Maksym Krutko, Christopher N. Mayhew, David T. Breault, James M. Wells, Jay Thiagarajah, Amy E. O’Connell, Olga Kovbasnjuk, Michael A. Helmrath

## Abstract

**Background and Aims:** *WNT2B* mutations result in Diarrhea-9 (DIAR9), a congenital diarrhea syndrome with an extreme phenotype and unique histological defects. Attempts to model DIAR9 in rodents and study patient epithelial tissue have not been able to fully reproduce the human phenotype, making understanding this condition challenging. Here, we aimed to interrogate the mechanisms and the specific cellular compartment contributing to DIAR9 using a human intestinal organoid model.

**Methods:** Human intestinal organoids (HIOs) generated from both a patient-derived WNT2B^R69*^iPSC line and a control line were transplanted into immunocompromised mice for 10 weeks. Grafts were harvested and histologically compared. Bulk RNA sequencing was performed on both organoid groups and on patient-biopsy derived enteroids. *In vitro* recombination experiments were performed to describe the causative cellular compartment.

**Results:** Live and histological imaging revealed partial epithelial delamination in WNT2B^R69*^HIOs, which was absent in controls. A significant number of crypts in WNT2B^R69*^ HIOs lacked OLFM4, a surrogate marker of stem cell activity. Key transcriptomic pathways altered between groups included trafficking of apical digestion proteins, which was confirmed via immunofluorescence. Patient derived enteroid proteomic analysis revealed similar results. Recombination experiments in HIOs revealed that while both epithelial and mesenchymal WNT2B are important for stem cell function, lack of mesenchymal WNT2B was sufficient to elicit the phenotype.

**Conclusion:** We demonstrated that mesenchymal WNT2B is critical for supporting human intestinal epithelial development and function.

## INTRODUCTION

Chronic diarrheal illnesses are a major threat to pediatric health, as they account for ∼7% of global childhood deaths (Zella and Israel, 2012). While most diarrheal diseases result from postnatal infections, some of the most severe cases fall under the umbrella category of congenital diarrhea and enteropathies (CODES) (Terrin et al., 2012; Thiagarajah et al., 2018). CODES patients have a poor quality of life, as they experience feeding intolerance, malabsorption, and fail to thrive, which requires extensive hospitalizations and parenteral nutrition for survival (Terrin et al., 2012; Thiagarajah et al., 2018). Prior to the advent of widespread genetic testing, the causes of CODES were largely unknown. A growing number of whole-genome studies on CODES patients have implicated a diverse set of monogenic mutations, the vast majority of which directly impair epithelial or immune cell function (Terrin et al., 2012; Thiagarajah et al., 2018; Ye et al., 2019). Understanding CODES may provide important insight into the mechanisms of enteral nutrient and fluid balance during homeostasis and following injury.

Recently, a subset of CODES patients with an extreme phenotype known as Diarrhea-9 (DIAR9) were identified (Tsukiyama and Yamaguchi, 2012; O’Connell et al., 2018; Zhang et al., 2021). These patients, who come from three distinct family backgrounds, presented with intractable, nutrient-independent osmotic diarrhea, growth failure, and years-long dependence on parenteral nutrition (O’Connell et al., 2018; Zhang et al., 2021). While the mechanism underlying DIAR9 was not identified, whole exome sequencing identified mutations in *WNT2B* as the root cause (O’Connell et al., 2018; Zhang et al., 2021). Histological evaluation of patient biopsies demonstrated aberrant epithelial architecture, including partial villous atrophy and paucity of crypts (O’Connell et al., 2018; Zhang et al., 2021). Understanding the mechanisms underlying these phenotypic defects will be critical for developing therapeutic strategies for these patients and may provide a broader understanding of chronic diarrhea in developing infants.

It has been demonstrated that Wnt signaling is essential for supporting both the embryonic development and adult homeostatic function of the intestine (Cervantes et al., 2009; Kondo and Kaestner, 2021). Globally disrupting Wnt signaling in both neonatal and adult mice results in aberrant crypt and villus morphology, loss of intestinal stem cells, and death within two weeks (Korinek et al., 1998; Valenta et al., 2016). Seven distinct Wnt proteins are expressed in the murine intestine; three are predominantly expressed by epithelial cells, and four, including Wnt2b, are classically thought to be expressed by mesenchymal cells (Gregorieff et al., 2005). Indeed, it was recently demonstrated that Wnt2b is specifically expressed by the murine telocytes, a fibroblast lineage thought to be important for maintain the stem cell compartment at the base of the crypt (Aoki et al., 2016; Shoshkes-Carmel et al., 2018). This expression pattern potentially makes the etiology of DIAR9 unique compared to other CODES disorders, which result from epithelial and immune cell dysfunction (Valenta et al., 2016; Miyoshi, 2017; Flanagan et al., 2018; Holloway et al., 2021). Existing data has recently called the source and phenotype of WNT2B loss into question. First, damage to epithelial-only human colonoids *in vitro* results in epithelial expression of *WNT2B* as part of the regenerative response (In et al., 2020). Moreover, establishing enteroids from the intestines of DIAR9 patients was remarkably inefficient, even with the addition of exogenous murine Wnt2b, also suggesting a potential role in epithelial WNT2B in regeneration or homeostasis (O’Connell et al., 2018). Determining whether the origin of the defect lies in epithelium or in mesenchyme is crucial to gaining insight into disease pathophysiology. We hypothesized that compartmental failure of both the epithelium and mesenchyme, to produce functional WNT2B caused pleiotropic impacts on the epithelium, leading to improper crypt and villus formation and regenerative dysfunction that manifests chronic diarrhea.

One major roadblock for studying DIAR9 *in vivo* is that the murine model does not phenocopy human disease. Multiple studies have found that knocking out *Wnt2b* in mice fails to generate an intestinal phenotype other than an increased susceptibility to DSS-induced colitis (van Amerongen and Berns, 2006; O’Connell et al., 2023). Additionally, studies on primary patient specimens are difficult, as availability is limited and biopsy materials do not represent full thickness tissue, inhibiting study of the mesenchyme. Moreover, enteroids from these patients are difficult to passage, propagating poorly. Human intestinal organoids derived from induced pluripotent stem cells (iPSCs) are an ideal system to overcome these constraints (Spence et al., 2011). While *in vitro* HIOs represent early first trimester fetal intestine, transplanting HIOs results in maturation of both the intestinal epithelium and mesenchyme, resulting in structures reminiscent of more mature tissue (Watson et al., 2014). Because of the cellular diversity within HIOs, they serve as a great tool for elucidating the mechanisms underlying DIAR9.

To test our hypothesis, we generated and transplanted HIOs from iPSCs that were derived in-house from a DIAR9 patient’s fibroblasts. After 10 weeks post-engraftment, the transplanted HIOs (tHIOs) were harvested, histologically examined, and compared to control tHIOs and to enteroids created from DIAR9 patient biopsies to confirm phenotype replication. Next, bulk RNA sequencing was performed to examine the key biological pathways that differed between both types of tHIOs and compared to the key biological pathways that differed between patient and control-derived enteroids. Enteroids were also derived from both types of tHIOs and used to assess functional differences between the types of grafts. Finally, to determine the compartmental role of WNT2B to the disease, a rescue experiment was performed in which the epithelium and mesenchyme of DIAR9 HIOs were separated from each other and reattached to the mesenchyme and epithelium of control HIOs, respectively, prior to transplant. Thus, we used tHIOs as a model system to gain insight into this novel intestinal disease.

## Materials and Methods

### Enteroid Derivation and Culture

Enteroids were derived from patient biopsies as previously described (Sato et al., 2011; Mahe et al., 2015). Briefly, the sample was digested with Collagenase Type 1, mechanically fragmented by pipetting, and incubated at 37°C. Next, samples were repeatedly pipetted to dissociate the crypts. The reaction was stopped with Advanced DMEM/F-12 (Thermo Fisher), and the samples were centrifuged at 500g for 5 minutes. The supernatant was aspirated, and the crypts were resuspended in Matrigel (BD Biosciences) prior to replating in 50 µL Matrigel bubbles. Media supplemented with rock inhibitor was provided every 2-3 days until enteroids formed. Once enteroids were ready to split, they were incubated in Cell Recovery Solution (Corning), centrifuged, and resuspended in fresh Matrigel. The same technique was applied to tHIOs to obtain the tHIO-derived enteroids used in this study, as previously described (Vales et al., 2020).

### WNT2B^-/-^ iPS Generation

Fibroblasts from a three-year-old male donor with DIAR9 were cultured in MEF media (DMEM, 10% FBS, 1mM non-essential amino acids). Cells at ∼50% confluency were transduced overnight with Sendai viral vectors (Cytotune 2.0, ThermoFisher Scientific) at MOIs of 2.5 (Klf4,Oct4,Sox2), 2.5 (cMyc), and 1.5 (Klf4). Fresh MEF media was provided on days 1,3, and 5 post-transduction. On day 7, transduced cells were plated in MEF media on irradiated MEF feeders (187,500 cells/well) in 6 well plates coated with 0.1% gelatin. On day 8, spent MEF media was removed and replaced with hESC media (DMEM/F12, 20% knockout Serum replacement, 1mM L-Glutamine, 0.1mM beta-mercaptoethanol, 1x non-essential amino acids, 2µg/mL bFGF). Starting on d8, wells underwent a complete daily media change with 2.5mL hESC media. Putative iPSC colonies were manually excised and replated in feeder free culture conditions consisting of Stem cell qualified Cultrex (BioTechne) and mTeSR1 (StemCell Technologies). Lines exhibiting robust proliferation and maintenance of stereotypical human pluripotent stem cell morphology were expanded and cryopreserved at passage 10.

### HIO Generation

*In vitro* HIOs were derived from the H1 embryonic stem cell line (WiCell Research Institute, Inc.) that was CRISPR modified to express GFP at the AAVS1 locus and the WNT2B^-/-^ iPS Line, as previously described (Spence et al., 2011; Poling et al., 2018). Briefly, stem cells were grown on Matrigel-coated plates and provided with mTESR1 media (Stem Cell Technologies). Colonies were dissociated with Accutase (STEMCELL Technologies), and cell numbers were quantified with a TC20 Automated Cell Counter (Bio-Rad) prior to plating at a density of 100,000 cells per well. Cells were grown in mTESR1 for two days, and 100 ng/ml of Activin A (Cell Guidance Systems) in Definitive Endoderm (DE) induction media (RPM1 1640, 100x NEAA, and 0.2-2% dFCS) for three days. Next, DE was furnished with hindgut induction media (RPMI 1640, 100x NEAA, 2% dFCS,) supplemented with 100 ng/ml FGF4 (R&D) and 3 µM Chiron 99021 (Tocris) to induce hindgut spheroids. After four days, spheroids were harvested, replated in Growth Factor Reduced Matrigel, and supplied with Minigut media (Advanced DMEM/F-12, N2 supplement, B27 supplement, 15 mM HEPES, 2 mM L-glutamine, penicillin-streptomycin) supplemented with 100 ng/ml EGF (R&D) to promote HIO maturation. Minigut media was changed biweekly. After 14 days, HIOs were individually cultured until transplantation at Day 28.

### Organoid Recombination

After 14 days, HIOs from both the H1-GFP and the WNT2B^-/-^ iPS Line were used for organoid recombination experiments. Briefly, HIOs were incubated in Cell Recovery Solution to dissolve the Matrigel as well as to depolymerize the basement membrane. After incubation, HIOs were gently pipetted to separate the epithelium from the mesenchyme, which detached as single cells. Mesenchymal cells were filtered with a 40-µm cell strainer (Fisher), counted with a TC20 Automated Cell Counter, and plated in 96-well Ultra Low Attachment Plates (Corning Costar) at a density of 50,000 cells per well. Intact pieces of epithelium were seeded onto the mesenchymal cells and cultured in a 1:1 mixture of Minigut media and IntestiCult Oragnoid Growth Medium (Stem Cell) overnight. The next day, recombined HIOs were individually replated in Matrigel and cultured in the same mixed media for three days. Minigut media was then provided on a biweekly basis until transplantation at Day 28.

### Animal Handling and HIO Transplantation

After 28 days *in vitro*, HIOs were transplanted into male and female nonobese diabetic, severe combined immunodeficiency, interleukin-2Rγnull (NSG) *mus muscularis* between the ages of eight and sixteen weeks, as previously described (Watson et al., 2014; Singh et al., 2020a; Singh et al., 2020b). Animal handling was performed in accordance with the NIH Guide for the Care and Use of Laboratory Animals. Mice were maintained on an antibiotic chow (275p.p.m. sulfamethoxazole and 1365p.p.m.trimethoprim; Test Diet) in the pathogen-free vivarium at Cincinnati Children’s Hospital Medical Center. Food and water were provided *ad libitum*. The Institutional Animal Care and Use Committee at CCHMC (Building and Rebuilding the Human Gut, Protocol No. 2021-0060) approved all animal experiments.

Each mouse was administered 2.5% inhaled isoflurane (Butler Schein) as an anesthetic. The left flank was shaved and sterilized with isopropyl alcohol and povidine-iodine. One cm incisions were made in the posterior subcostal skin and retroperitoneal muscle. A single HIO was implanted into the renal subcapsular space (RSS). The kidney was replaced, and the peritoneal cavity was flushed with a 2.38 mM solution of piperacillin-tazobactam (AuroMedics). Incisions were closed using a double layer closure technique, and mice were injected with carprofen (1 mg/mL) for pain management.

### Tissue Harvest

After 8-10 weeks, mice were humanely euthanized, and grafts were harvested. A portion of each graft was flash frozen in liquid nitrogen for RNA sequencing, and a portion was fixed for immunohistochemistry or RNAScope. Prior to fixation, the graft was imaged under a Leica M165 FC dissection microscope with a DCF7000 T camera (Leica Microsystems) to assess for epithelial integrity. Five grafts were harvested per group.

### Tissue processing, Immunohistochemistry, Immunofluorescence and Microscopy

Half of each tHIO was fixed using either 4% paraformaldehyde overnight or formalin for five days. After fixation, samples were processed, embedded in paraffin, and sectioned. The staining protocol has been outlined previously (Poling et al., 2018; Singh et al., 2020b). All antibody incubation steps were performed overnight at 4ºC. Dilutions and references for markers assessed are listed in Table 1. Stain images were captured with a Nikon Eclipse Ti and analyzed using Nikon Elements Imaging Software (Nikon).

### RNA Scope

In situ hybridization on formalin-fixed paraffin sections was conducted with the RNAscope Multiplex Fluorescent Reagent Kit v2 and manual assay RNAscope catalog probes (Advanced Cell Diagnostics). Probe hybridization was detected using Opal Dye 520 and 570 fluorophores (Akoya Biosciences) following the respective vendor-supplied protocols. Slides were counterstained with Hoechst 33342 and coverslipped with ProLong Gold Antifade mountant. Fluorescent images were acquired on an Olympus FV3000 scanning laser confocal microscope and processed in ImageJ2/FIJI.

### RNA Isolation and Sequencing

An RNAeasy Plus Micro Kit (Qiagen) was used to extract RNA. RNA concentration was quantified using a NanoDrop One (Thermo Scientific) and submitted for Next Generation Sequencing to the DNA Sequencing and Genotyping Core. After quality control, a cDNA library was generated, and sequencing was performed using an IlluminaHiSeq2000 (Illumina) at a depth of 20 million paired-end reads per sample.

### RNA Sequencing Processing and Analysis

The read quality of the fastq datasets was assessed using FastQC (v0.12.1) and MultiQC (v1.14). Reads were aligned to human genome assembly hg38 and quantified using the quasi-mapper kallisto (v0.46.1) with default settings. Differential gene expression analysis was performed using R package DESeq2 (v1.38.3). Size factors were utilized to normalize the read count matrix, and a variance stabilizing transformation (VST) was applied to the normalized expression data. Principal component analysis (PCA) was performed using the stats (v4.2.1) package. The gene with the highest loadings for each principal component was extracted from the PCA and plotted using the hi_loadings function in the pcaExplorer (v2.22.0) package. Functional enrichment analysis was performed in the ToppGene suite (https://toppgene.cchmc.org). The data was visualized using ggplot2 (v3.4.0), clusterProfiler (v4.4.4), GOplot (v1.0.2) and pcaExplorer (v2.22.0) packages.

### Protein Extractions

Total protein was extracted from enteroids using 2% SDS and probe tip sonication. Sample proteolysis, isobaric mass tag labeling, peptide fractionation, and liquid chromatography-tandem mass spectrometry was conducted by the Mass Spectrometry & Proteomics Core, Johns Hopkins University School of Medicine.

### Proteomics Data Processing

Tandem MS/MS spectra were searched using Mascot (version 2.8.0, Matrix Science) against the database RefSeq 2021_204_Human and processed in Proteome Discoverer (version 2.4, ThermoFisher) to identify peptides with ≤1% false discovery rate and to calculate protein ratios between groups. Results were filtered for p≤0.05 and a ratio fold change of 1.5 for inclusion in subsequent analysis.

### Proteomics Data Analysis

Protein accession identifiers were converted to gene symbols using the SynGO ID mapping tool (https://syngoportal.org/convert). GO-term enrichment was assessed via the DAVID Functional Annotation Tool (Huang da et al., 2009; Sherman et al., 2022). A list of genes derived from the calculated protein abundance ratios (WNT2B mutant versus wild-type enteroids, p-value ≤ 0.05) were compared to the background Homo sapiens to determine enrichment (Benjamini adjusted p-value ≤ 0.05). All other settings were default.

### Enteroid monolayer culture

Permeable tissue culture inserts (0.4 µm pore Transwells, 3470, Corning) were coated with human collagen type IV (10 µg/cm2, C5533, Sigma) in 100 mM acetic acid. Enteroids were fragmented by repeated aspiration with a P200 pipette, washed with DMEM, centrifuged at 300g, and resuspended in NDM media (Advanced DMEM/F12, 50% L-Wnt3a conditioned media, 15% HA-Rspondin1-Fc conditioned media, 10% Noggin-Fc conditioned media, 1x B27, 1 mM N-acetylcysteine, 50 ng/mL EGF, 100 µg/mL Primocin) containing 10 µM Y-27632 and CHIR 99021. The fragment suspension was plated in the upper compartment of the collagen-coated inserts. The lower compartment was supplied with media and plates were incubated at 37 °C, 5% CO2. Culture media was changed every 2-3 days. Inserts were inspected daily to determine progress toward confluence.

### Transepithelial Resistance

Transepithelial electrical resistance (TEER) in enteroid monolayers was measured using a voltohmmeter equipped with chopstick electrode probe (EVOM2, World Precision Instruments) and readings were corrected for surface area (0.33 cm2).

### Fatty acid uptake and staining

Enteroid monolayers were incubated in a chemically defined lipid mixture (1:100 in DMEM/F-12, Thermo 11905) for one hour. Monolayers were washed with PBS, fixed with 4% paraformaldehyde, treated with 60% isopropanol,s and stained with Oil Red O solution. Imaging was done with a Zeiss AxioObserver light microscope.

### Forskolin-induced swelling assay for CFTR activity

CFTR function in three-dimensional transplant-derived enteroid cultures was compared using the previously described forskolin-induced swelling assay (Dekkers et al., 2013). Enteroids were plated in Matrigel and cultured for 3 days in expansion medium. Calcein red-orange (1 μM, ThermoFisher C34851) was added to the media to label cells 1 h prior to initiating the assay. The plate was transferred to an environmentally-controlled (37 °C, 5% CO2) microscope chamber (Keyence BZ-X710) and 4 unique regions of interest were mapped for each condition in each independent experiment. Forskolin (10 μM) was added at t=0 to stimulate CFTR activity, and mapped regions were imaged under brightfield and fluorescence every 10 min over 1 h. Enteroid cross-sectional areas in the time-lapse fluorescent images were measured in ImageJ2/Fiji.

### Data Representation, Statistics and Reproducibility

Five grafts were harvested per group across three distinct batches, based on historical data about number of grafts required to validate differences between groups, and the number of grafts used was adjust on an per assay basis. For IHC, IF, RNAScope, morphometric analysis: five grafts were assay were used. For morphometric analysis, only grades 3 and 4 lumens were analyzed, and a minimum of 10 crypts and villi were measured per graft. Areas with active delamination were excluded from morphometric analysis. For analysis of OLFM4 expression, all crypts present in the graft (minimum of 10) were analyzed. For RNA sequencing, proteomics, and enteroid monolayer analysis: three grafts or enteroid bubbles were sequenced per group. All statistical data was represented as violin plots with a dashed line indicated the mean and interquartile ranges, and the individual data points represented. For statistics comparing the WNT2B^R69*^ tHIOs to the healthy control tHIOs: a student t-test was performed with the cutoff for statistical significance set at below 0.05.

## RESULTS

We previously reported difficulty with propagating WNT2B^R69*^ patient derived enteroids, (Sato et al., 2009) even upon supplementation of conditioned growth media with recombinant murine Wnt2b (O’Connell et al., 2018). This obstacle, coupled with the fact that enteroids only contain the epithelial compartment of the human intestine, led us to pursue a developmental model that contains both epithelium and mesenchyme to provide a novel opportunity to study this unique human disease. We thus created a WNT2B^R69*^ iPS line (Fig. S2) and opted to use these to generate HIOs (Fig. S1) to further study how loss of functional WNT2B leads to intestinal pathologies and diarrheal disease.

After 28 days *in vitro*, both WNT2B^R69*^ and control HIOs were further matured via transplantation under the kidney capsule of immunocompromised mice. tHIOs were harvested at eight to ten weeks post-transplantation, histologically examined, and compared to images of both control human duodenum and duodenum from a WNT2B^R69*^/DIAR9 patient. Unlike the control (Fig. 1A), the WNT2B^R69*^ patient’s duodenum displayed areas in which the epithelium were detached from the underlying basement membrane (Fig. 1B). Similarly, in comparison to control tHIOs and healthy human tissue (Fig. 1C), the WNT2B^R69*^ tHIOs (Fig. 1D) exhibited numerous regions of epithelial delamination. To determine whether this finding was a histological artifact, we examined grafts under live microscopy. Unlike control tHIO epithelium (Fig. 1E), WNT2B^R69*^ tHIO epithelium was consistently observed as free-floating and partially detached from the underlying tissue (Fig. 1F). Moreover, the areas with intact epithelium appeared to display aberrant mucosal morphology, which was also observed when we re-examined intestinal biopsy samples from a patient with WNT2B^R69*^. Specifically, WNT2B^R69*^ demonstrated significant reductions in crypt depth (Fig. S3A), villus height (Fig. S3B), and percent crypt fission (Fig. S3C) than controls. While we did observe aberrant morphology in control patient tissue, we did not morphometrically characterize this due lack of histological material. Thus, WNT2B^R69*^ tHIOs recapitulated major gross pathologies seen in intestinal tissue from WNT2BR69*/DIAR9 patients.

**Figure 1.**
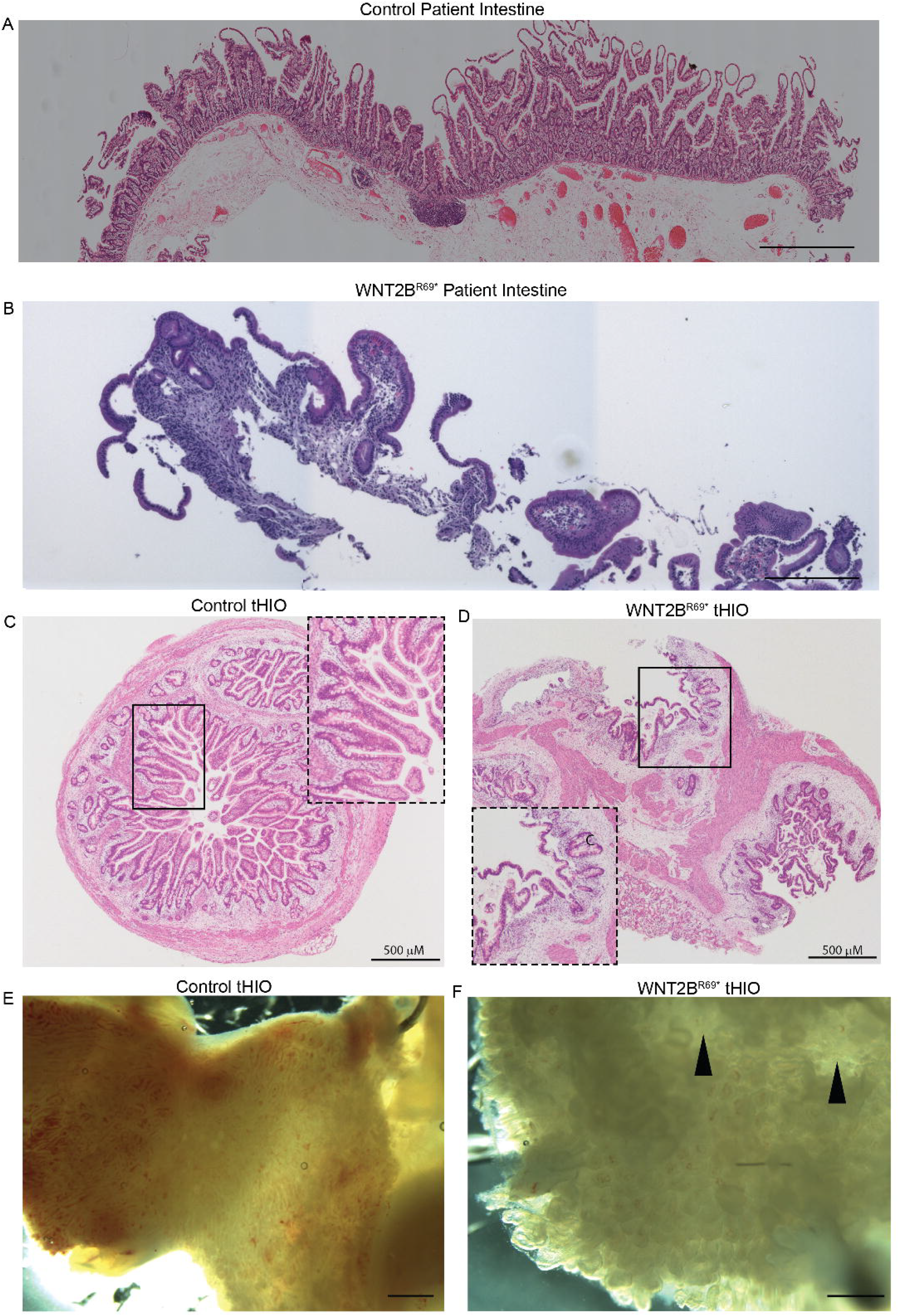
WNT2B^R69*^ mutation results in compromise of epithelial layer integrity. Representative images of H&E staining of **(A)** a surgical sample from a healthy patient, **(B)** a biopsy of a WNT2B^R69*^ patient **(C)** a control tHIO and **(D)** a WNT2B^R69*^ tHIO. Representative live imaging of **(E)** control tHIO and **(F)** WNT2B^R69*^ tHIO. Arrowheads indicate areas of epithelial delamination.

Previous findings suggested defects in intestinal stem cell function, as WNT2B^R69*^ patients’ intestine exhibit attenuated expression of OLFM4 (O’Connell et al., 2018) and overall crypt loss. We investigated if this finding was consistent in our model system using immunohistochemistry (IHC). We found a significant reduction in the percentage of crypts expressing OLFM4 in WNT2B^R69*^ tHIOs (Fig. 2A-2B). To further characterize the etiology of this in tHIOs, we examined mRNA expression of intestinal stem cell marker Leucine-rich repeat-containing G-protein coupled receptor 5 (*LGR5*) (Barker et al., 2007) and proliferation marker Cyclin Dependent Kinase 1 (*CDK1*). RNAScope revealed significant reduction in both the number of LGR5+ and CDK1+ cells in WNT2B^R69*^ tHIOs (Fig. 2C-2D, Fig. S3D). Finally, staining for Marker of Proliferation Ki-67 (MKI67) demonstrated restriction to the crypts in control tHIOs, but more diffuse expression in WNT2B^R69*^ tHIOs (Fig. 2E).

**Figure 2.**
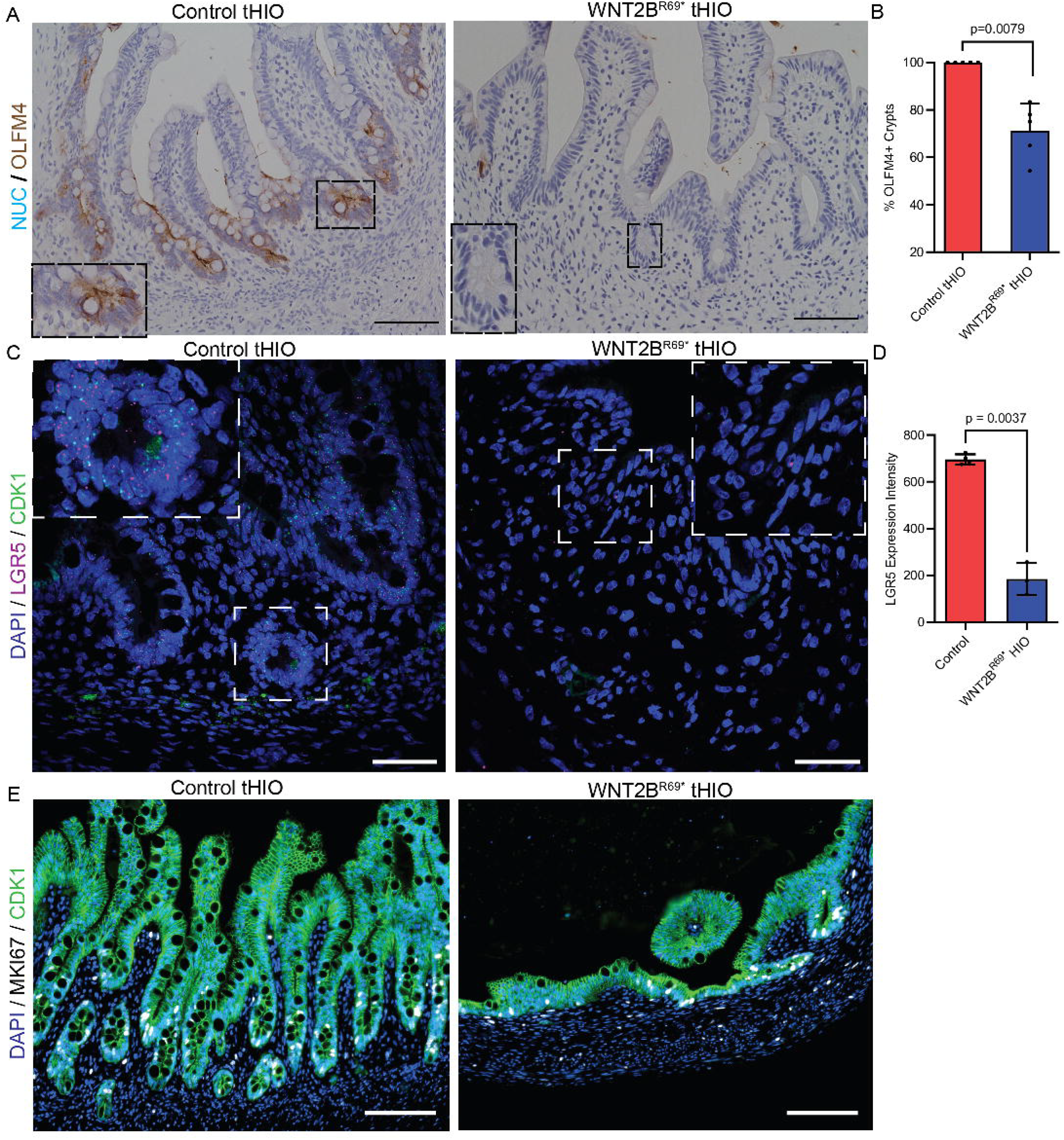
WNT2B^R69*^ tHIOs have an aberrant stem cell compartment. **(A)** Immunhistochemistry for OLFM4 (brown), a surrogate marker of stem cell activity **(B)** Bar graphs of the percentage of OLFM4+ crypts in each graft type. **(C)** RNAScope for stem cell marker LGR5 (purple) and proliferation marker CDK1 (green). **(D)** Bar graphs of LGR5 intensity in each graft type. **(E)** Immunofluorescence for proliferation marker MKi67 (white) and CDH1 (green) in each graft type.

We also examined differentiated secretory cell lineages in WNT2B^R69*^ tHIOs. Staining for Paneth cell marker Lysozyme (LYZ) revealed aberrant localization of Paneth cells outside of intestinal crypts in WNT2B^R69*^ tHIOs (Fig. S4A). Staining for goblet cell marker Mucin 2 (MUC2), and enteroendocrine cell marker Chromogranin A (CHGA) revealed similar expression patterns in both groups of tHIOs (Fig. S4A-43B). However, 5-hydroxytryptamine (5HT), a marker of enterochromaffin-like cells, which secrete 5HT to regulate gut motility, (Gross et al., 2016), was missing entirely from WNT2B^R69*^ tHIOs (Fig. S4C).

To gain insight into the signaling pathways altered in WNT2B^R69*^ tHIOs, we performed bulk RNA Sequencing on both types of grafts and on control and WNT2B^R69*^ patient-derived enteroids. Both WNT2B^R69*^ tHIOs (Fig. 3A) and enteroids (Fig. S5A) clustered separately from their respective controls, suggesting disease related transcriptomic alterations. Some of the key downregulated Biological Processes in WNT2B^R69*^ tHIOs included “epithelial tube development”, “angiogenesis”, “negative regulation of wound healing”, and “response to nutrient level” (Fig. 3B). Similar key biological processes were downregulated in WNT2B^R69*^ enteroids (Fig. S5B), with the addition of “digestion” and “lipid transport”. Moreover, in both WNT2B^R69*^ tHIOs and enteroids, alterations were found in genes impacting cellular compartments (Fig. 3C, Fig. S5C). Specifically, genes found in the “apical part of the cell”, “brush border”, and “microvillus membrane” were downregulated in the disease state. Finally, assessment of altered molecular function revealed downregulation of pathways related to nutrient transport (Fig. 3D, Fig. S5D), including “lipid transport”, “carbohydrate binding”, and “receptor ligand activity”.

**Figure 3.**
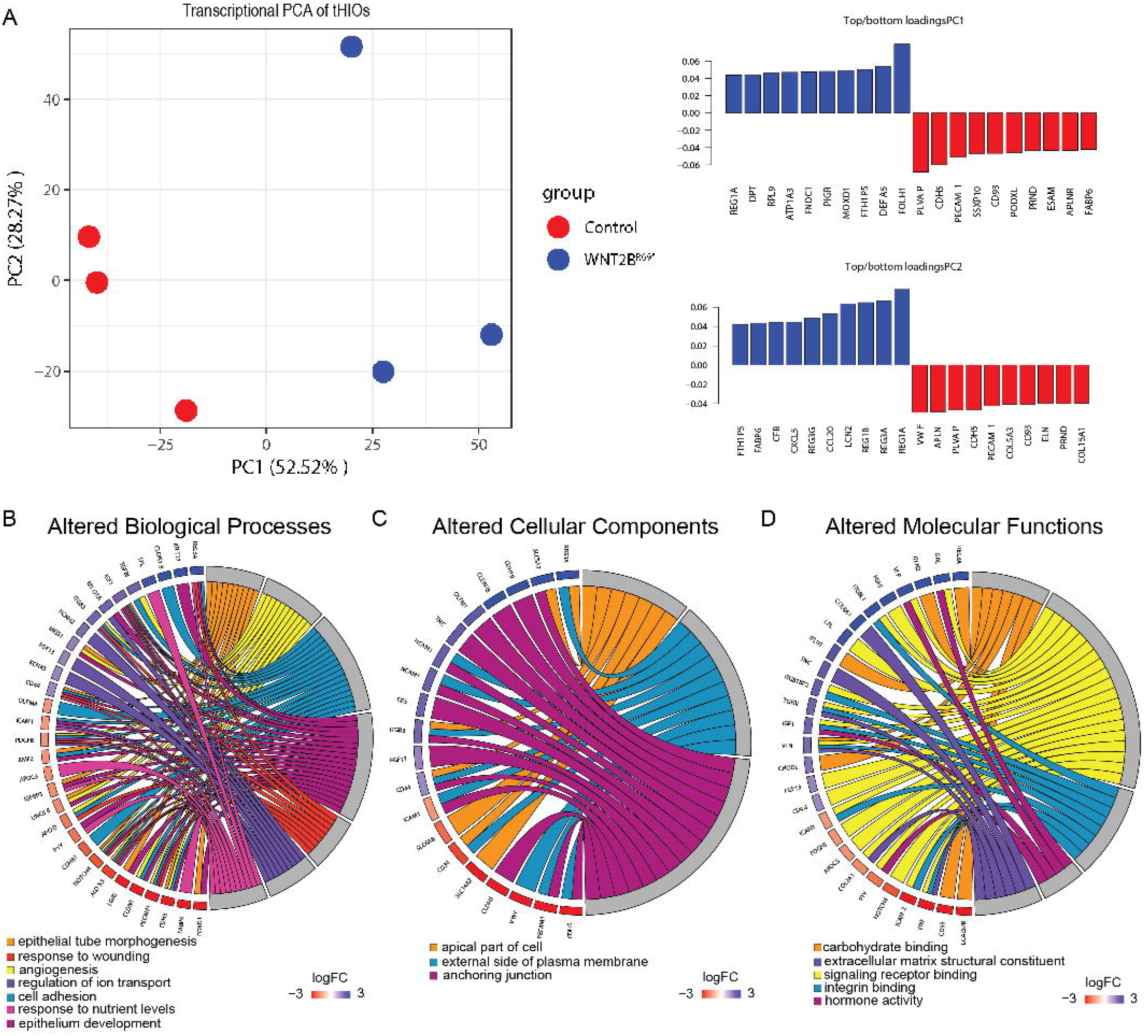
Key intestinal pathways are altered because of mutations in *WNT2B*. **(A)** PCA plot of WNT2B^R69*^ and control tHIOs reveals transcriptional separation between graft types. Key altered **(B)** biological pathways, **(C)** cellular components, and **(D)** molecular functions in WNT2B^R69*^ tHIOs (red) as compared to controls (blue).

Based on these results, as well as the fact that restricting any given nutrient failed to prevent diarrhea in the patients (O’Connell et al., 2018), we next examined tHIOs for expression of key apical and basolateral proteins involved in digestion and cell anchoring. Sucrase-Isomaltase (SI), which is involved in carbohydrate digestion, was missing entirely from WNT2B^R69*^ tHIOs, but not in controls (Fig. 4A). Similarly, Fatty Acid Binding Protein 2 (FABP2), which is involved in the absorption of lipids, had reduced expression in WNT2B^R69*^ tHIOs, but not in controls (Fig. 4B). The expression of Laminin Subunit Alpha 1 (LAMA1), a key protein involved in anchoring the epithelium to the basement membrane (Timpl et al., 1979; Teller et al., 2007), appeared to be absent in areas of epithelium that were delaminating from the underlying mesenchyme (Fig. 4C) while expressed in control tissue. However, the expression of Laminin Subunit Gamma 1 (LAMC1) was found to be the same in both types of tHIOs (Fig. 4D).

**Figure 4.**
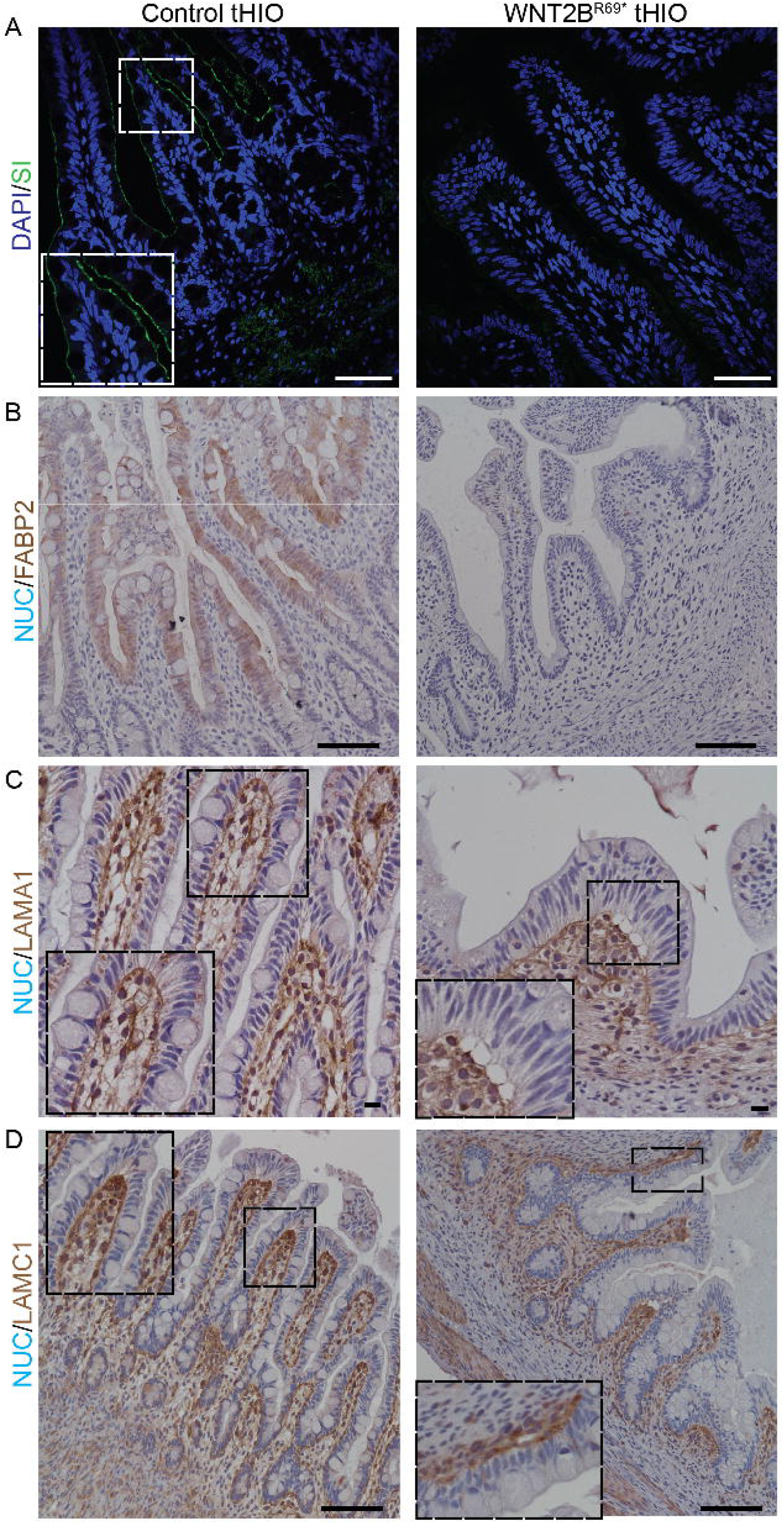
WNT2B^R69*^ tHIOs have defects in key apical and basolateral proteins. Immunofluorescence for **(A)** carbohydrate digestive enzyme SI (green). **(B)** Immunohistochemistry for fatty acid transporter FABP2 (brown) in both types of grafts. **(C)** Immunohistochemistry for basolateral protein LAMA1 (brown) in both types of grafts. **(D)** Immunohistochemistry for basolateral protein LAMC1 (brown) in both types of grafts.

One major advantage of enteroids is the ability to conduct robust, *in vitro* functional experiments. We thus sought to determine whether WNT2B^R69*^ tHIOs could be used as a source of crypts/stem cells for enteroid culture. tHIO derived Enteroids (it-Enteroids) were successfully derived from both control and WNT2B^R69*^ tHIOs (Fig. S6A) and passaged over time in Intesticult media. We further used the it-Enteroids for functional assessment of the cystic fibrosis transmembrane conductance regulator protein (CFTR), which can be activated via exposure to forskolin (Boj et al., 2017). it-Enteroids from WNT2B^R69*^ tHIOs did not swell as much as controls (Fig 5A, Fig. S6B). Additionally, we generated monolayer cultures, which are widely used for nutrient transport assays (Sanman et al., 2020). While assessing nutrient transport deficits, we found that WNT2B^R69*^ derived monolayers had difficulty absorbing lipids, as assessed through staining for Oil Red O (Fig. 5B). Finally, we used the monolayer culture system to study differences in the trans-epithelial resistance (TEER) between the WNT2B^R69*^ it-Enteroids and the control it-Enteroids. At each measured timepoint, the WNT2B^R69*^ it-Enteroids had a higher TER than the control it-Enteroids (Fig 5C).

**Figure 5.**
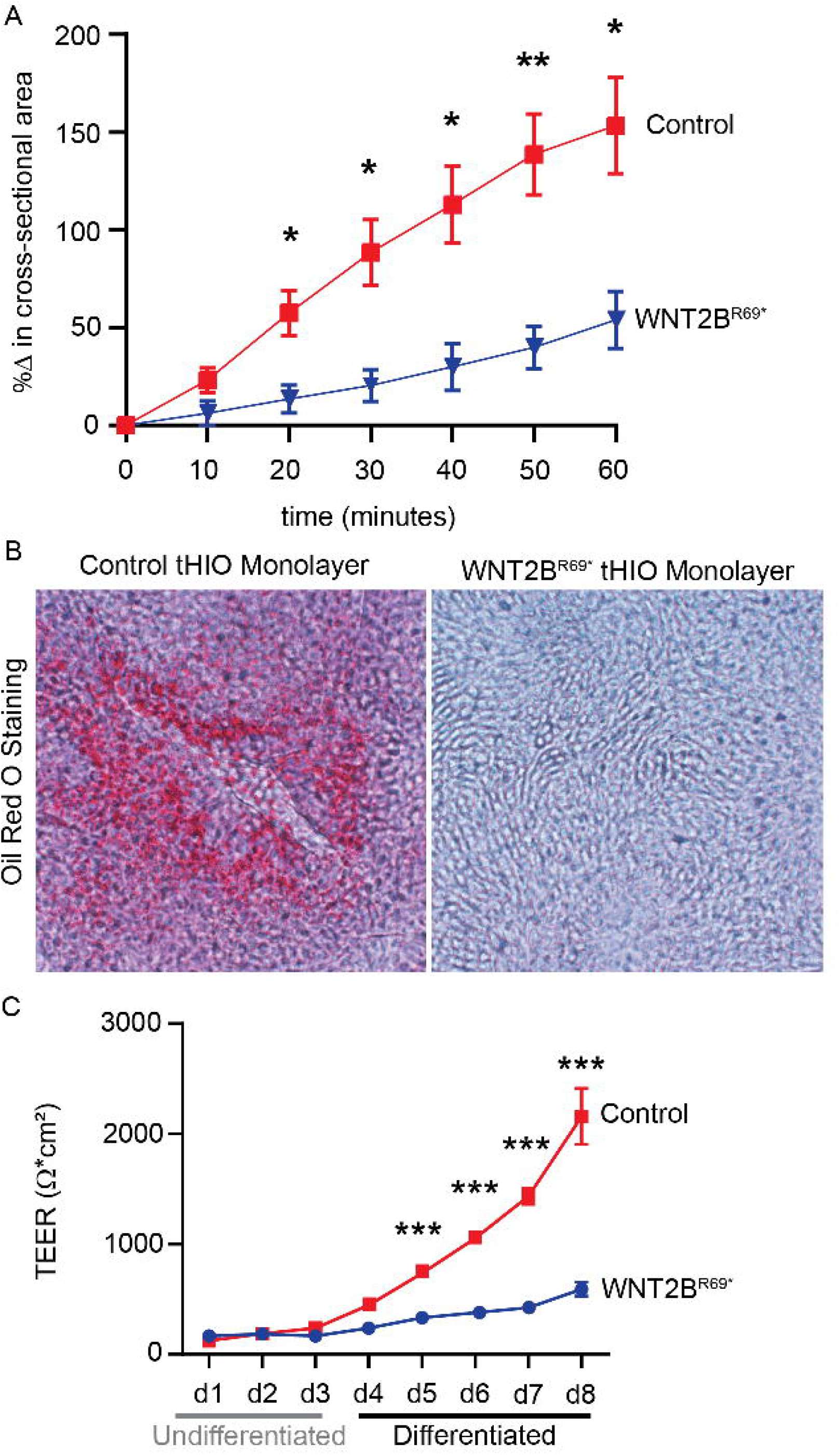
Enteroid cultures derived from WNT2B^R69*^ tHIOs reveal functional epithelial deficits. **(A)** Results of forskolin assay on enteroids from both graft types **(B)** Representative images of Oil Red O staining after uptake study in monolayers from both graft types. **(C)** TEER measurements across time from monolayers derived from both graft types.

To further interrogate alterations in protein expression in the epithelium, we extracted total protein from WNT2B^R69*^ and control it-Enteroids for mass spectrometry. Analysis of the proteomics data revealed clustering of the samples in accordance with disease status (Fig. 6A-B). Some of the notable pathways that were downregulated in the WNT2B^R69*^ it-Enteroids included “protein transport”, “maintenance of epithelial apical/basal polarity”, “positive regulation of integrin-mediated signaling pathway”, and “positive regulation of cell adhesion”, which were similar to the transcriptional pathways found to be altered by RNA sequencing (Fig. 6C). Importantly, expression of LAMA1 was found to be downregulated in WNT2B^R69*^ it-Enteroids.

**Figure 6.**
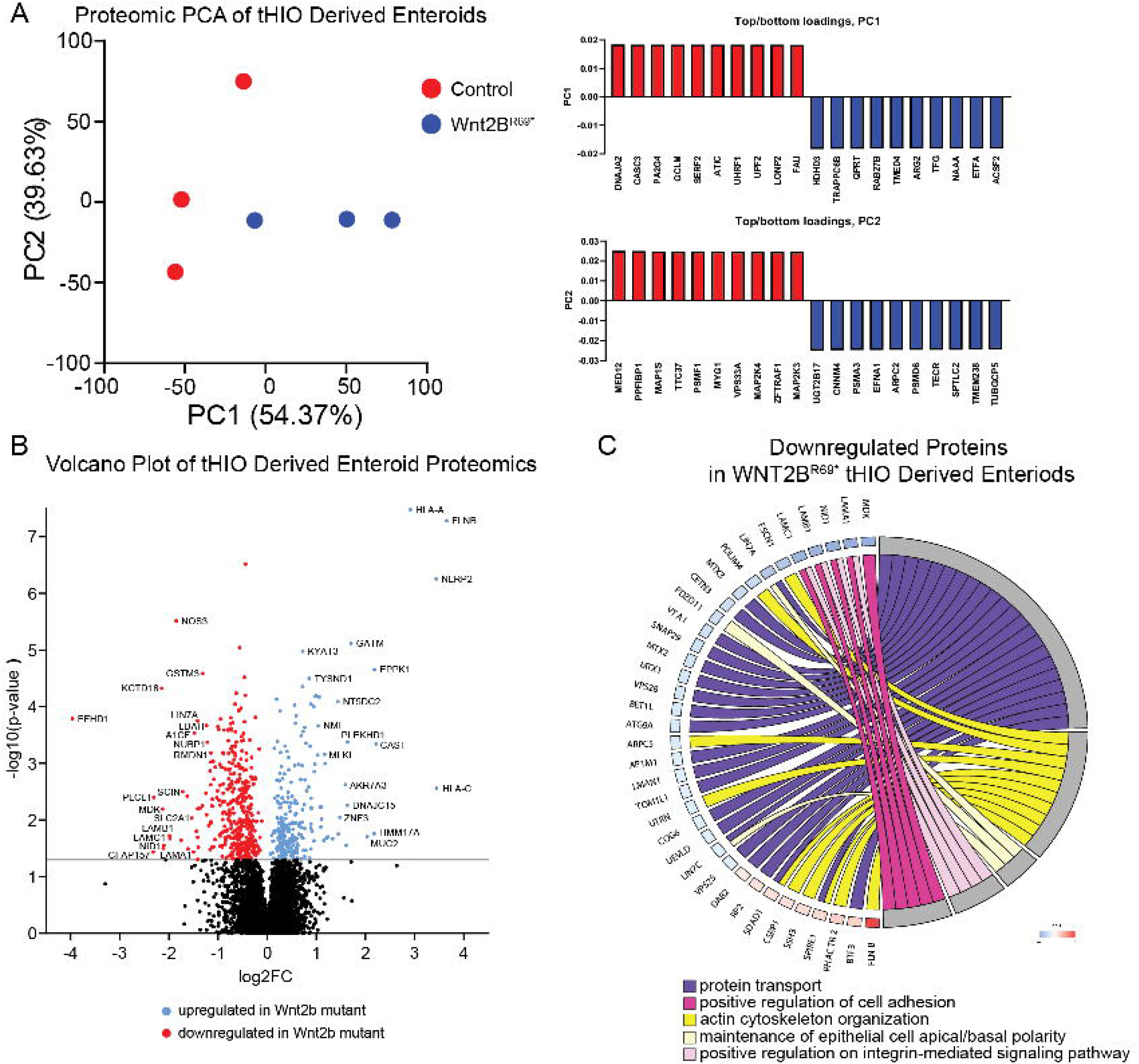
Proteomic analysis of enteroids derived from both graft types reveals alterations in cell adhesion and apical/basolateral proteins. **(A)** PCA of proteomic data from enteroids reveals separation based on graft type. **(B)** Volcano plot of differentially expressed proteins between the two groups of enteroids. **(C)** Circos diagram of key biological processes downregulated at the protein level in WNT2B^R69*^ enteroids.

Finally, to determine the compartmental contribution to the phenotype, we performed a series of epithelial and mesenchymal dissociations of HIOs from both groups followed by recombinations and transplantation (Fig. 7A). This served as a test to determine whether healthy, or control, mesenchyme could rescue the WNT2BR69* epithelial phenotype, or if the WNT2BR69* epithelium alone was sufficient to cause the phenotype. In all, four distinct groups of HIOs were created through recombination: Endo^Cont^/Meso^Cont^, Endo^R69*^ / Meso^R69*^, Endo^R69*^ /Meso^Cont^, and Endo^Cont^/ Meso^R69*^ (Fig. S7). HIOs from each group were transplanted into immunocompromised mice, harvested after 8-10 weeks, and histologically examined. Unsurprisingly, tHIOs from the Endo^Cont^/Meso^Cont^ group displayed an intact crypt/villus axis, and those from the Endo^R69*^ / Meso^R69*^ group showed aberrant development of the epithelium, with areas of the pseudostratified epithelium characteristic of early gut development (Fig. 7B). Moreover, portions of the epithelium in the Endo^R69*^ / Meso^R69*^ group appeared to be delaminating from the basement membrane. Additionally, as expected, staining for OLFM4 revealed robust expression throughout the crypts in the Endo^Cont^/Meso^Cont^ group, and heterogeneous expression in the Endo^R69*^ / Meso^R69*^ group (Fig. 7C, 7E). Finally, staining for LYZ revealed localization to the base of the crypts in the Endo^Cont^/Meso^Cont^ group, but ectopic expression in the Endo^R69*^ / Meso^R69*^ group. (Fig. 7D).

**Figure 7.**
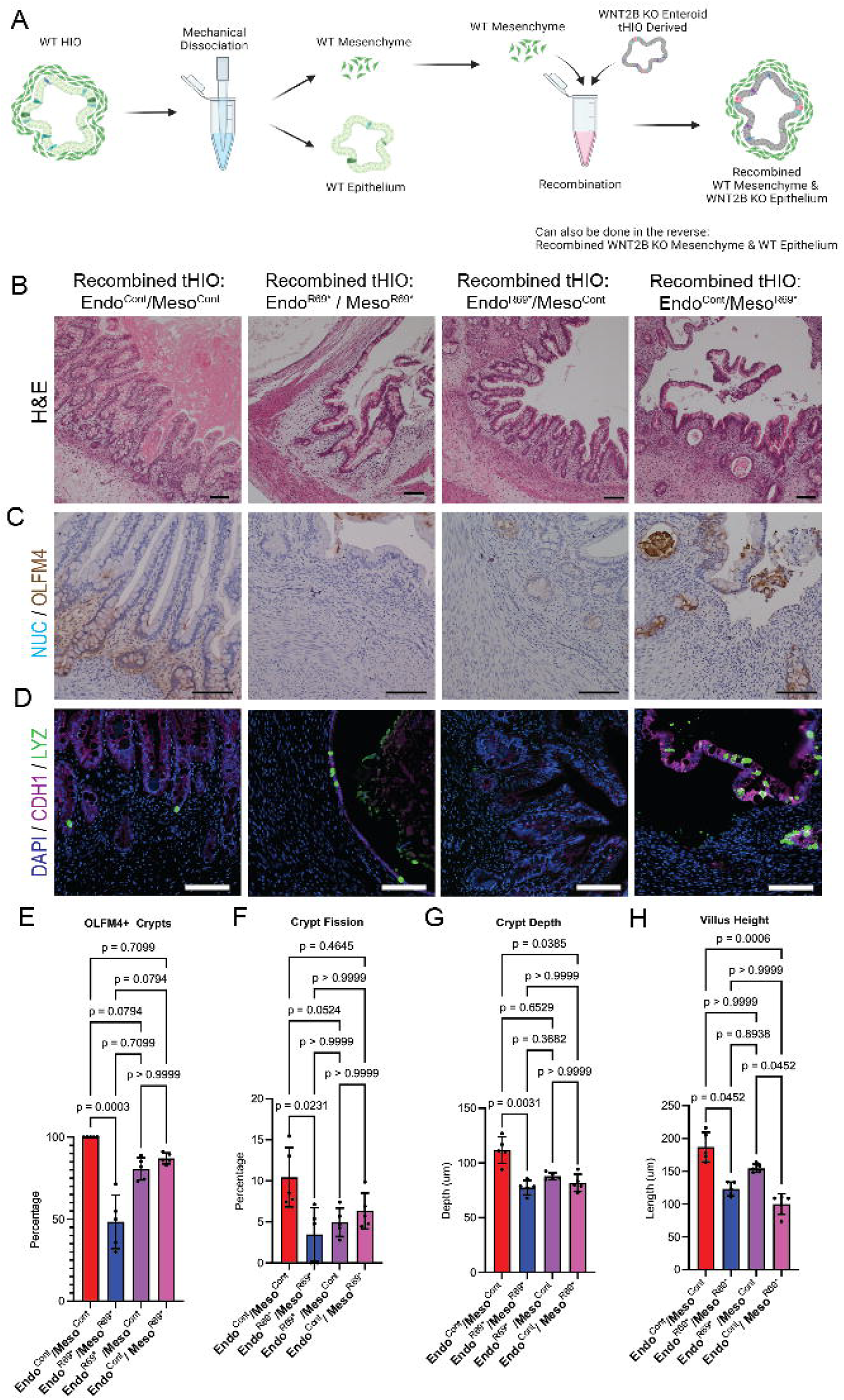
Recombination experiment reveals that while both the epithelial and mesenchymal compartments contribute to the defect, lack of mesenchymal WNT2B is sufficient to generate the phenotype. **(A)** Schematic outlining the recombination procedure **(B)** Representative images of H&E stains of tHIOs reveals delamination of the epithelium in groups with WNT2B^R69*^ mesenchyme. **(C)** Staining for OLFM4 (brown) in tHIOs from each recombination group. **(D)** Staining for CDH1 (purple) and LYZ (green) in tHIOs from each recombination group. **(E)** Bar chart of the percentage of OLFM4+ crypts in grafts of each type. **(F)** Bar chart of precent crypt fission, **(G)** crypt depth, and **(H)** villus length in each group.

In contrast to the tHIOs in the Endo^R69*^ / Meso^R69*^ group, the tHIOs in the Endo^R69*^ /Meso^Cont^ group showed an intact developed crypt/villus axis. This was not the case with the tHIOs in the Endo^Cont^/ Meso^R69*^ group, which demonstrated some areas with less developed epithelium and others in which the epithelium detached from the basement membrane (Fig. 7B). Staining for OLFM4 revealed heterogenous expression in both groups (Fig. 7C, 7E). Staining for LYZ showed proper expression in the Endo^R69*^ /Meso^Cont^ groups, but ectopic expression in the Endo^Cont^/ Meso^R69*^ group (Fig. 7D). Percent crypt fission was only significantly lower than the Endo^Cont^/Meso^Cont^ group in the Endo^R69*^ / Meso^R69*^group (Fig. 7F). Crypt depth and villus height, however, were significantly lower in the Endo^R69*^ / Meso^R69*^and Endo^Cont^/ Meso^R69*^ groups, but not the Endo^R69*^ /Meso^Cont^ group (Fig. 7G-H).

## DISCUSSION

In comparison to all other model systems, human intestinal organoids have the unique capacity to model human patient-specific intestinal diseases in an *in vivo* setting (Singh, 2020; Workman, 2017; Krishnamurthy, 2022). Because they can be generated from iPSCs, they can be used to develop an understanding of a disease at the level of an individual patient (Zhang et al., 2019; Krishnamurthy et al., 2022), truly allowing for the realization of personalized medicine. While incredible insight into intestinal development has been gleaned from animal models, new insights into human intestinal development will benefit from *in vivo* human models that are experimentally tractable (Singh et. al, 2023). Thus, tHIOs are an excellent tool for studying intestinal development and disease.

While it was clinically recognized that DIAR9 patients presented at the time of birth with an osmotic diarrhea without a specific deficit in absorption of any specific nutrient (O’Connell et al., 2018; Zhang et al., 2021), the mechanisms underlying the disease pathogenesis remained elusive. Although Wnt2b is clearly expressed by the intestinal telocytes, studying this disorder with a murine model is difficult given that the murine phenotype is more subtle, only leading to increased susceptibility to DSS-induced colitis (Aoki et al., 2016; Miyoshi, 2017; Shoshkes-Carmel et al., 2018; O’Connell et al., 2024). Given that both telocyte ablation as well as abrogation of Wnt expression by telocytes leads to destruction of crypt/villus architecture in the murine intestine, it is likely that other telocyte Wnts are able to compensate for the loss of Wnt2b (Aoki et al., 2016; Shoshkes-Carmel et al., 2018). Indeed, it has been previously suggested that Wnt2b and Wnt3 may act in a redundant manner in the murine intestine, as genetic deletion of either protein individually fails to nullify stem cell activity *in vivo*, and Wnt2b is able to rescue stem cell activity in enteroid cultures that lack Wnt3 in an *in vitro* setting (Farin et al., 2012). Given the observed phenotype in patients with *WNT2B* mutations, it is likely that either this redundancy is not preserved in the human intestine, or that a different, currently unknown Wnt protein in the murine intestine is analogous to WNT2B in the human intestine. In either case, a human model of the condition provides further insight.

Our data suggests that the chronic diarrhea that these patients suffer from stems from defects in the production of key hormones and digestive proteins, including those involved in carbohydrate and lipid absorption. Indeed, deficiencies in apical protein expression have previously been implicated in several CODES diseases, including microvillus inclusion disease (Overeem et al., 2016; Schlegel et al., 2018; Thiagarajah et al., 2018). While many of these other disorders result from monogenic mutations in specific transporters, *WNT2B*^R69*^ appears to result in a malabsorptive phenotype for multiple nutrient types.

In addition to defects in apical protein expression, our results indicated that there were defects in the basolateral portion of the cell, specifically in LAMA1, which is involved in attachment of the epithelial cells to the basement membrane. Previous work has suggested that basement membrane proteins are involved in establishing intestinal barrier function, as well as in influencing growth of the intestinal epithelium (Vllasaliu et al., 2014). Thus, loss of LAMA1 expression in these portions of the epithelium likely contributes to the diarrhea seen in these patients. Interestingly, previous work has shown that as an individual ages, LAMA1 expression in the gut is replaced by expression of Laminin subunit alpha-2 (LAMA2); LAMA1 expression is missing entirely from adult human intestine (Teller et al., 2007). This could potentially explain why the delamination phenomenon disappears in patients over time, as well as why the disease phenotype becomes less severe as the patients age.

One interesting finding regarding the epithelial delamination that was observed is that the while the cells detached from the underlying stromal population, they did not detach as individual cells, but rather remained tightly adherent to each other in sheets. Our RNA sequencing data suggests that this likely stems from the upregulation of cell-cell adhesion molecules, such as (Protocadherin Beta 6) *PCDHB6* and Protocadherin 20 (*PCDH20)*. This upregulation of cell-cell adhesion molecules likely explains why, in the monolayer system WNT2B^R69*^ monolayers exhibited a significantly higher TER than the controls once the cells were placed in a differentiation media. Indeed, no deficits in lateral cell-adhesion proteins, such as Epithelial-cadherin (CDH1), were ever observed in our system.

Although *WNT2B* has been thought to be expressed exclusively by the mesenchyme, recent evidence suggests that there is some *WNT2B* expression in the intestinal epithelium at baseline, and that it can be upregulated in injury conditions (In et al., 2020; Zhang et al., 2021; Xie et al., 2022). Our recombination study suggests that ultimately both epithelial and mesenchymal WNT2B are required for proper development and function of the intestinal epithelium, albeit with lack of WNT2B from the mesenchyme being sufficient to elicit the dominant disease phenotype. This suggests that the disease phenotype directly correlates with mesenchyme’s mutation status, and that mesenchymal WNT2B is important both for human intestinal stem cell function, formation of normal intestinal architecture, attachment of the epithelium to the mesenchyme, and apical digestive protein trafficking. Mesenchymal WNT2B is thus important for controlling the development of the epithelium functionally, developmentally, and spatially.

The phenotype of epithelial WNT2B^R69*^, however, is more subtle. Providing epithelium from a WNT2B^R69*^ HIO with the mesenchyme of a control HIO restored normal intestinal architecture, but did not completely restore intestinal stem cell function, as Endo^R69*^ /Meso^Cont^ tHIOs still showed mosaic expression of OLFM4. Providing the epithelium of a control HIO with the mesenchyme of a WNT2B^R69*^ HIO recapitulated the disease phenotype, with mosaic OLFM4 expression, abnormal crypt/villus architecture, and delamination of the epithelium from the underlying mesenchyme. This validates the emerging notion in the literature that epithelial WNT2B is important for human intestinal stem cell function.

One limitation of our findings is that tHIOs lack a functional immune system, and some inflammation was observed in the patients early in the disease course (O’Connell et al., 2018). Additionally, tHIOs were grown in a sterile environment, and were not exposed to gut flora or nutrients, while the intestines of these children have been exposed to these potential disease triggers. Furthermore, the tHIOs in this study lacked an enteric nervous system, which is important for regulating intestinal motility, barrier function, and nutrient absorption. We recognize these complexities were not recapitulated in our system and likely contribute to additional aspects of the disease. Despite these limitations, however, we are confident that the conclusions in our study can help inform patient care in the clinic. We were able to replicate the disease phenotype, and indeed deepen understanding of disease pathogenesis, suggesting that the tHIO system is an appropriate vehicle for studying this disorder. Moreover, the fact that this occurred in the absence of an immune system indicates that the inflammation results from, rather than is a driver of, disease pathogenesis. Indeed, our RNA sequencing results indicated that multiple genes associated with inflammation, leukocyte migration, and chemokine activity were upregulated in WNT2B^R69*^ tHIOs, suggesting that the inflammation is a byproduct of the intestinal damage in this disorder. Finally, we attempted to understand some of these differences by performing RNA sequencing on enteroids derived from the patient biopsies and found that similar pathways were altered.

Another limitation of our study is the use of a control group that was not genetically matched to the patient-derived WNT2BR69* iPSC line. Although our initial aim was to generate an isogenic control line by correcting the mutation, technical challenges prevented the successful creation of this line. As an alternative, we employed the well-characterized H1-GFP ESC line for comparative analyses. While this provided a reliable reference point, we acknowledge that it does not serve as the ideal isogenic control. Future mechanistic studies will benefit from the successful generation of a genetically corrected line to enable more precise comparisons.

Applying these findings to clinical care requires further study into disease pathogenesis. Because it appears that severity of disease varies on a case-by-case basis (O’Connell et al., 2018; Zhang et al., 2021), modeling and studying the differences between more severe and less severe phenotypes might allow for elucidation of therapeutic targets and strategies for these patients. Luckily, the tHIO model system is well suited for such a personalized approach and has been successfully used for this application in the past (Zhang et al., 2019). Indeed, it appears that over time, some of the patients can be weaned off parenteral nutrition, and that the disease phenotype ultimately abates as the patients age. Identification of the niche factors that compensate for lack of WNT2B over time could allow for the development of targeted medical therapies for these patients earlier in the course of the disease, potentially allowing clinicians to inhibit some of the more consequential impacts of the disease, such as failure to thrive. Additionally, it is possible that genetically corrected HIOs could serve as a therapeutic tool for healing the epithelial damage experienced by these patients as a source of cell therapy (Poling *et al*., 2024). This approach would have the advantage of avoiding transplant rejection, as well as providing a native mesenchymal source of WNT2B.

## Acknowledgments

The human iPSCs used in this study were generated by the Pluripotent Stem Cell Facility at Cincinnati Children’s Hospital Medical Center. The Veterinary Services facility at CCHMC supported the animal work in this project. The authors also thank the DNA Sequencing and Genotype Core at CCHMC for processing and sequencing the bulk RNA Sequencing data, as well as by the Mass Spectrometry & Proteomics Core at Johns Hopkins University School of Medicine for processing and sequencing the proteomics data. Schematics in Supplemental Figure 1 and 7A were created with Biorender.com Created in BioRender. poling, h. (2025) https://BioRender.com/z0g5gbu.

