## Supplemental Information for "Mesenchymal WNT2B is required for the development and function of the human intestine"

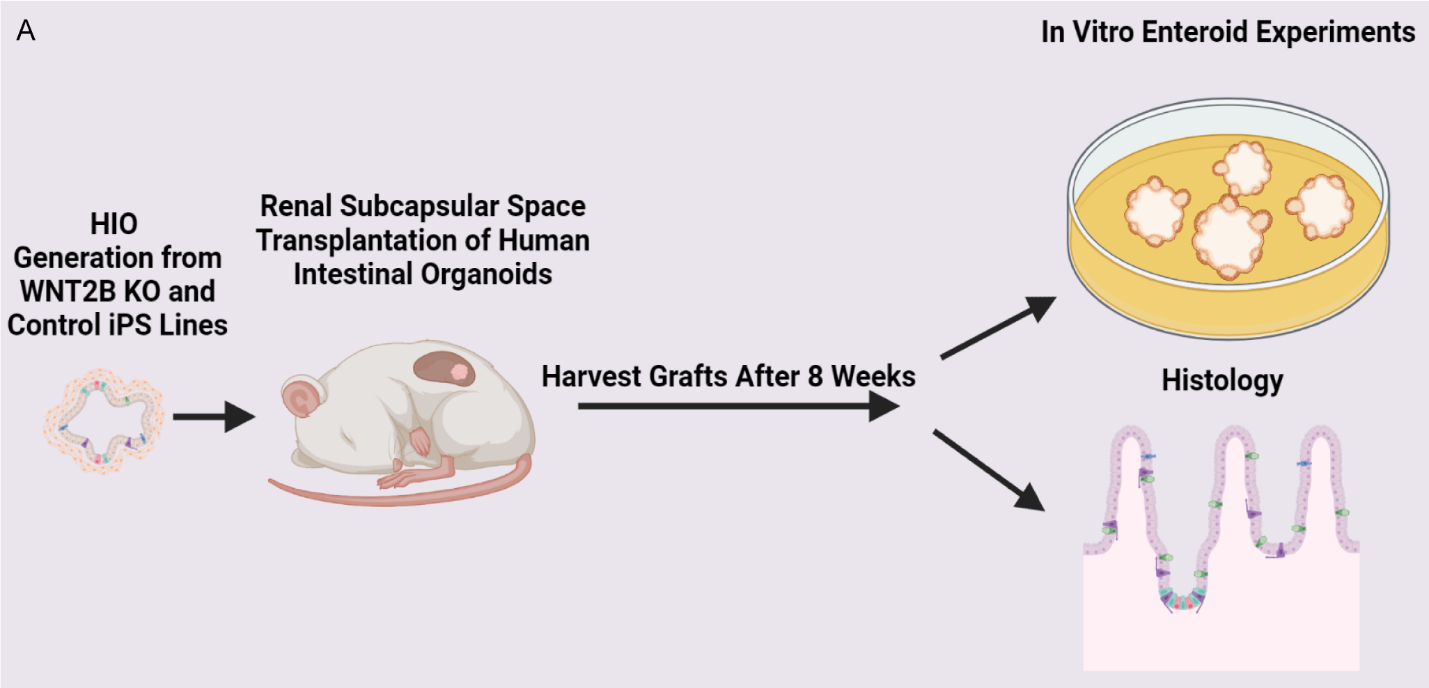

**Supplemental Figure 1: Graphical abstract of the experimental design**.

**­**
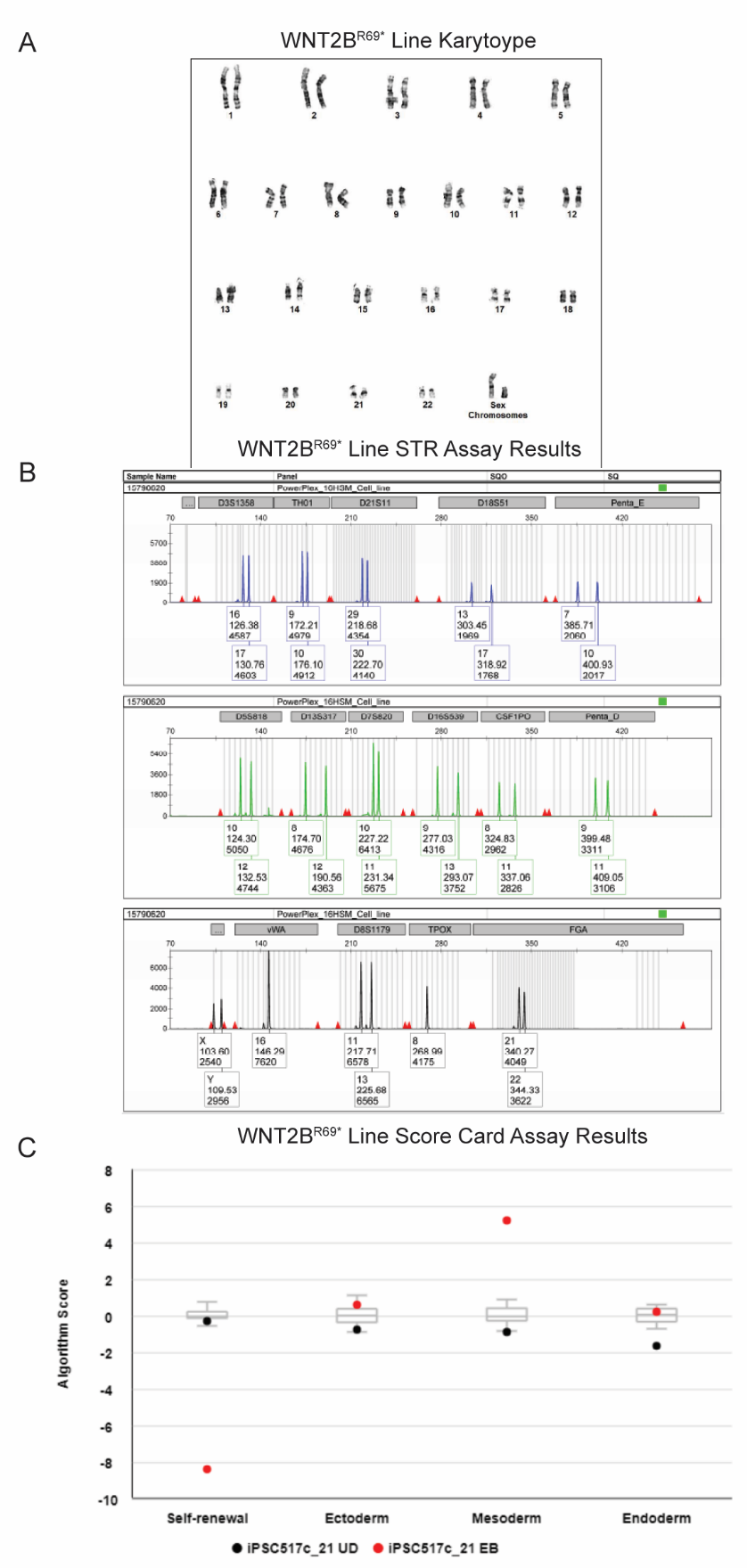

**Supplemental Figure 2: Quality improvement assays on the iPS line derived from a WNT2B^R69*^. (A)** Karyotype of the WNT2B^R69*^ iPS line. **(B)** STR assay of the WNT2B^R69*^ iPS line. **(C)** Scorecard assay of the WNT2B^R69*^ iPS line.

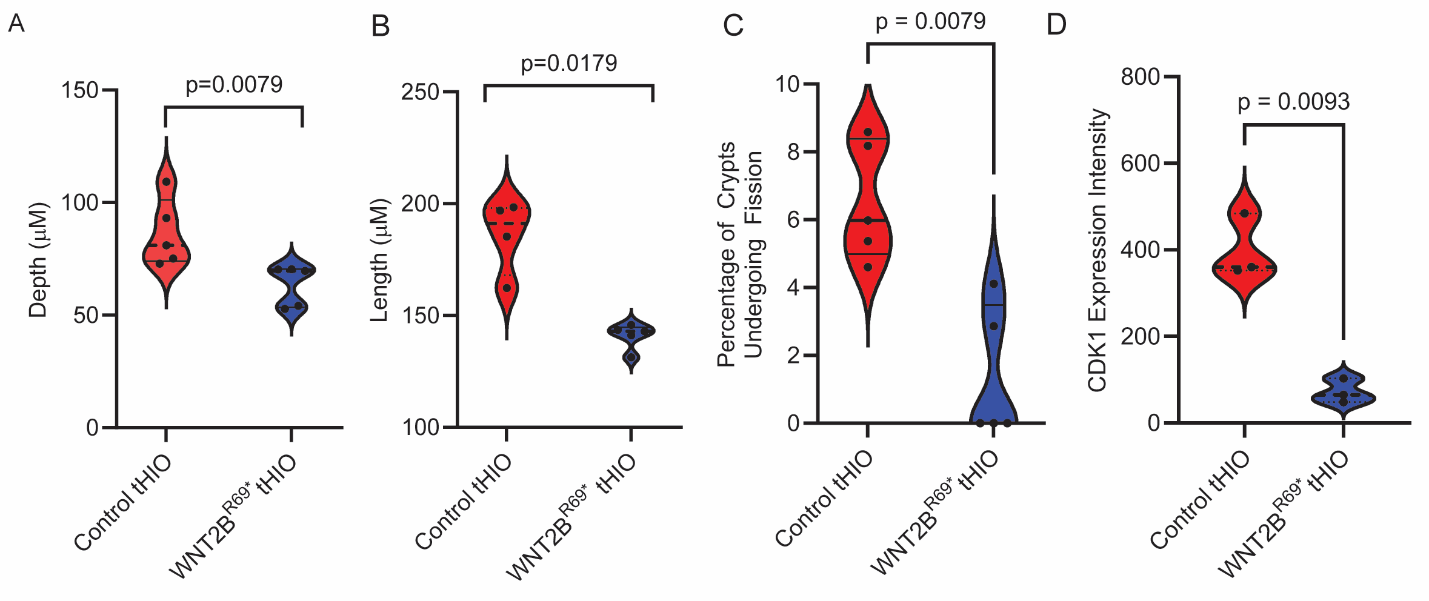

**Supplemental Figure 3: Epithelium of WNT2B^R69*^ tHIOs displays abnormal morphometric properties. (A)** Violin plots of the average crypt depth that is present in grafts of each type. **(B)** Violin plots of the average villus height that is present in grafts of each type. **(C)** Violin plots of the percent crypt fission that is present in grafts of each type. **(D)** Violin plots of CDK1 intensity in grafts of each type.

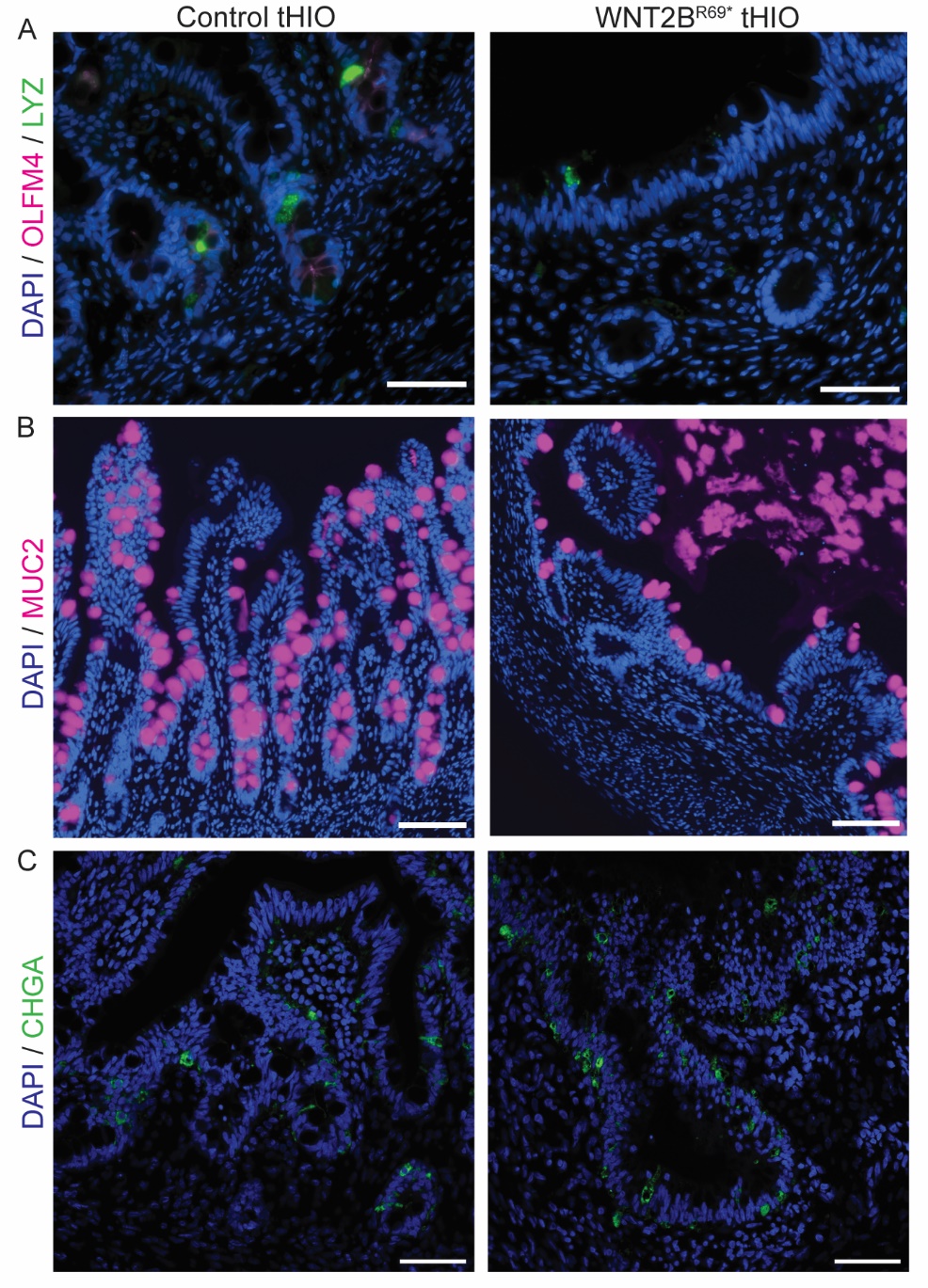

**Supplemental Figure 4: Staining for secretory epithelial cell types reveals lack of serotonergic cells in WNT2B^R69*^ tHIOs.** (A) Immunofluorescence for OLFM4 (pink) and Paneth cell marker LYZ (green) in each graft type. Immunofluorescence for (B) goblet cell marker MUC2 (pink). (C) enteroendocrine cell marker CHGA (green) in grafts of each type.

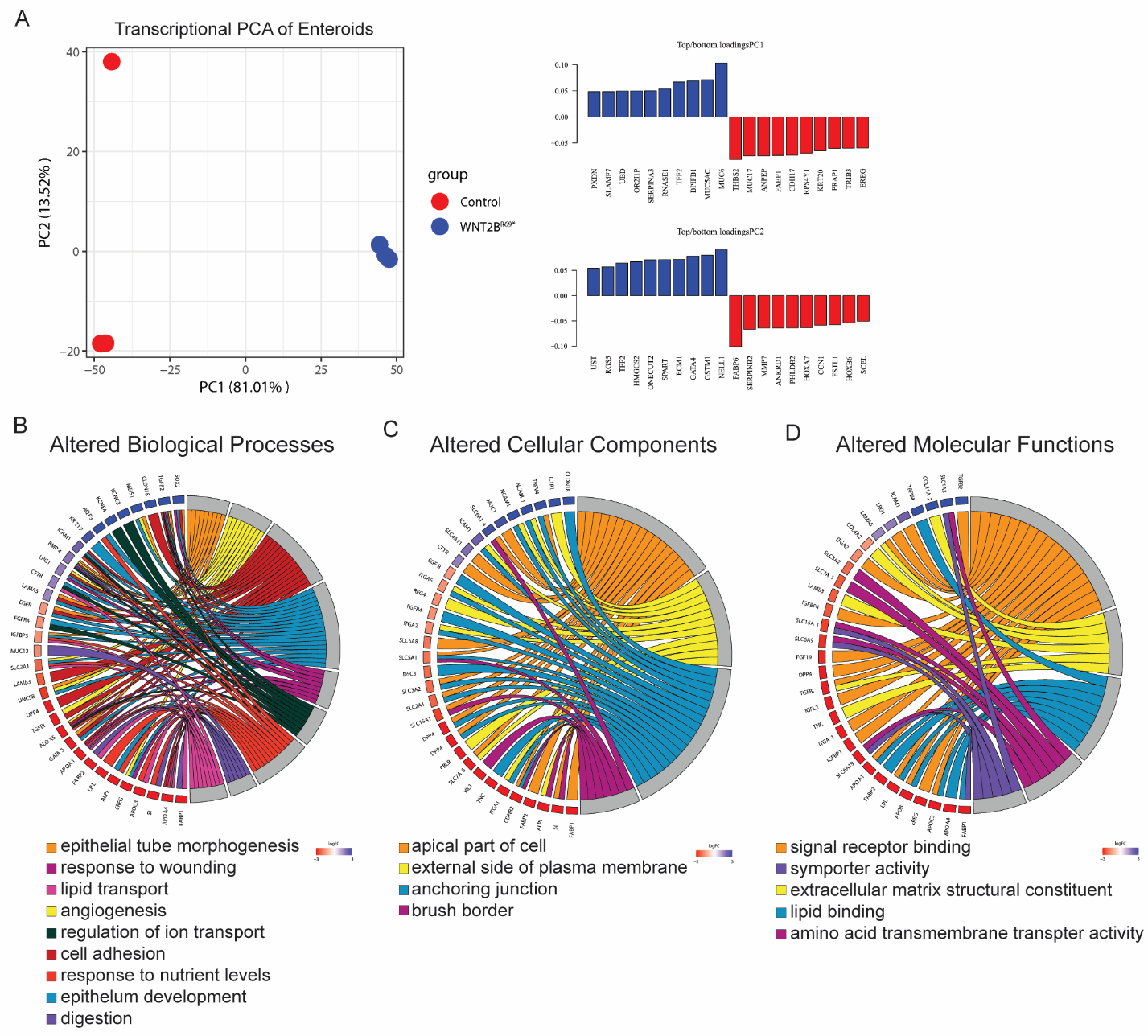

**Supplemental Figure 5: Key intestinal pathways are altered as a result of WNT2B^R69*^mutations.** (A) PCA plot of WNT2B^R69*^ and control enteroids reveals transcriptional separation along the lines of disease status (B) Key altered biological pathways in WNT2B^R69*^ patient-derived enteroids (red) as compared to controls (blue). (C) Key altered cellular components in WNT2B^R69*^ patient-derived enteroids (red) as compared to controls (blue). (D) Key altered molecular functions in WNT2B^R69*^ patient-derived enteroids (red) as compared to controls (blue).

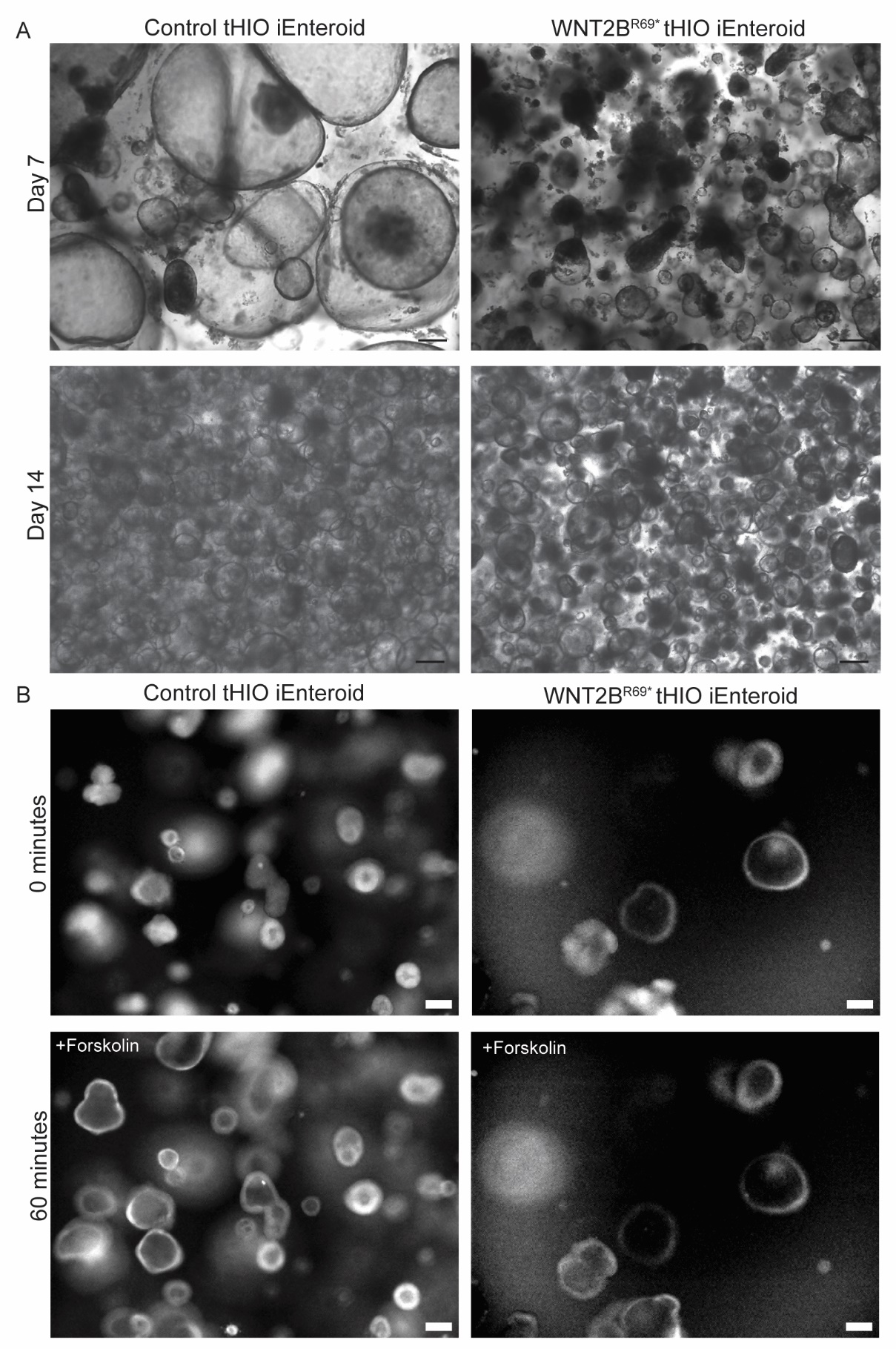

**Supplemental Figure 6: Enteroids can be derived from the crypts of WNT2B^R69*^ tHIOs and used for functional studies.** (A) Representative images of enteroids derived from control tHIOs and WNT2B^R69*^ tHIOs grown in high WNT media. (B) Representative images of enteroids derived from both types of grafts when exposed to forskolin for an hour.

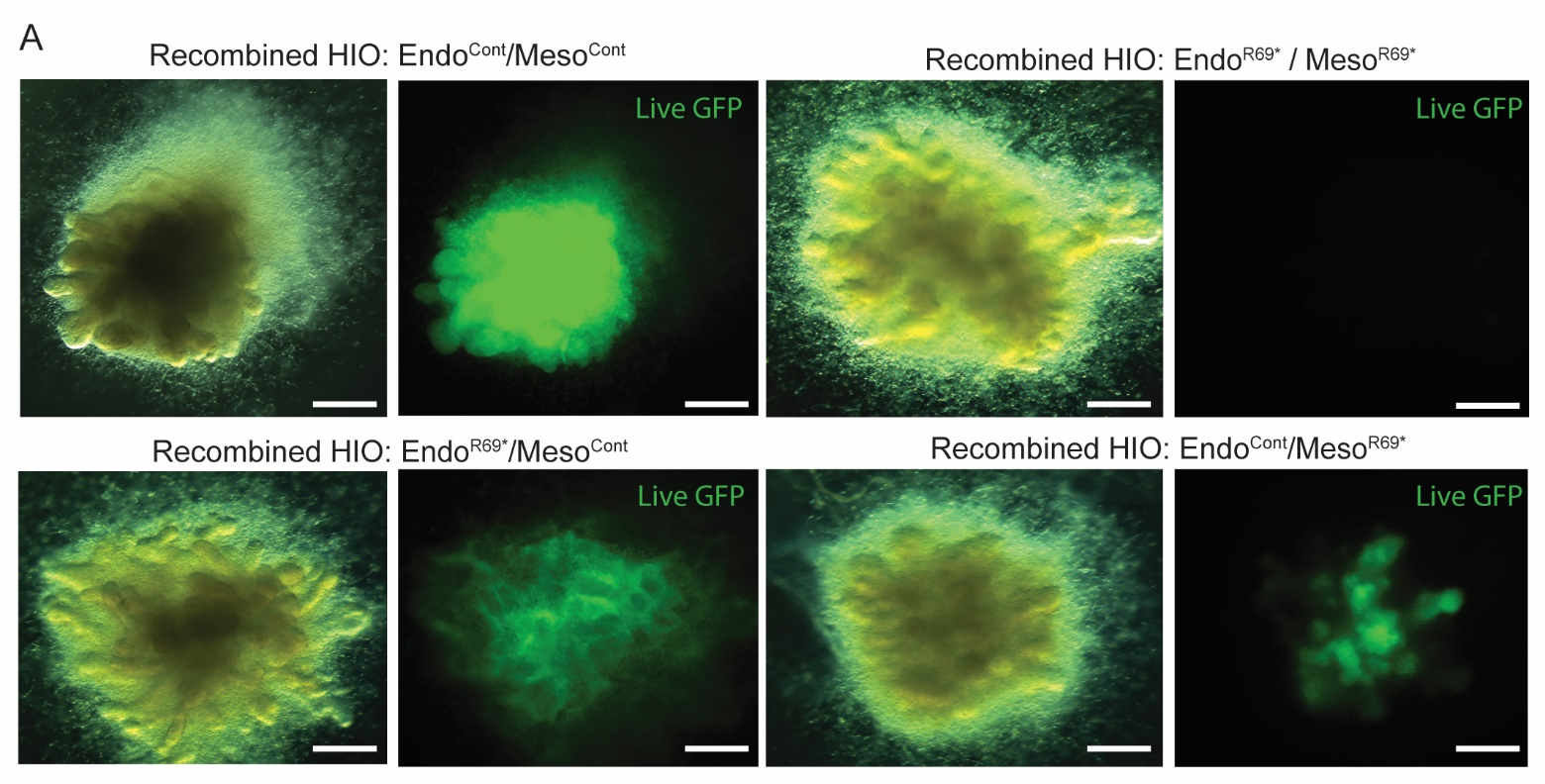

**Supplemental Figure 7: Recombination technique was used to successfully create *in vitro* HIOs with mixed compartmental origins.** Live imaging of representative *in vitro* HIOs from each recombination group. Cells from control HIOs expressed Green Fluorescent Protein.

|  | **Antigen** | **Dilution** | **Host** | **Company: Catalog Number** |
| --- | --- | --- | --- | --- |
| **Primary** | Serotonin (5HT) | 1:100 | Goat | Abcam: ab66047 |
|  | Chromogranin A (CHGA) | 1:100 | Mouse | Thermo Fisher: MA5-13096 |
|  | Defensin Alpha 5 (DEFA5) | 1:500 | Mouse | Abcam: ab90802 |
|  | E-Cadherin (CDH1) | 1:300 | Mouse | BD: 610182 |
|  | Fatty Acid Binding Protein 2 (FABP2) | 1:600 | Rabbit | Atlas: HPA034607 |
|  | Catenin Beta 1 (CTNNB1) | 1:100 | Mouse | BD Biosciences: 610154 |
|  | Lysozyme (hLYZ) | 1:2000 | Rabbit | Biorad: 0100-0523 |
|  | Marker Of Proliferation Ki-67 (MKI67) | 1:350 | Rabbit | Thermo Fisher: RM-9106-50 |
|  | Mucin-2 (MUC2) | 1:1100 | Rabbit | Abcam: ab134119 |
|  | Olfactomedin 4 (OLFM4) | 1:200 | Mouse | Cell Signaling: 14369 |
|  | Laminin Subunit Alpha 1 (LAMA1) | 1:500 | Rabbit | Novus: NB300-144 |
|  | Laminin Subunit Gamma 1 (LAMC1) | 1:500 | Rabbit | Novus: NBP1-87718 |
|  | Sucrase-Isomaltase (SI) | 1:800 | Rabbit | Sigma: HPA011897 |
| **Secondary** | α-rabbit AF647 | 1: 800 | Donkey | Life Technologies: A31573 |
|  | α-rabbit AF488 | 1: 800 | Donkey | Life Technologies: A21206 |
|  | α-rabbit Biotin | 1:1000 | Goat | Vector: BA-1000 |
|  | SA Cy3 | 1:500 |  | Abcam: ab175704 |
|  | α-goat AF647 | 1:800 | Donkey | Life Technologies:A21447 |
|  | α-mouse AF568 | 1: 800 | Donkey | Life Technologies: A10037 |

**Table 1: Antibody information for immunohistochemistry and immunofluorescence staining.**
